# The complete plastid genomes of two Fabaceae orphan crops from Africa

**DOI:** 10.1101/459248

**Authors:** Xuezhu Liao, Yang Liu, Huan Liu

**Affiliations:** BGI-Shenzhen, Shenzhen 518083, China; China National GeneBank, BGI-Shenzhen, Shenzhen 518120, China

**Keywords:** Orphan crops, plastid genome, phylogenetic analysis

## Abstract

Dolichos bean (*Lablab purpureus*) and Bambara groundnut (*Vigna subterranea*) are two traditional crops from Africa with important economic values. The study of such neglected or underused crops (orphan crops) should contribute to the resolution of food shortage in Africa. Here we assembled and described the complete plastid genome of the two Fabaceae orphan crop species. The length of the complete plastomes of *L. purpureus* and *V. subterranea* are 151,753 bp and 152,015 bp, respectively. Maximum-likelihood (ML) phylogenetic analyses indicated that *L. purpureus* and *Phaseolus vulgaris* are closely related, and *V. subterranea* and other *Vigna* species are clustered in one clade, which is congruent with former studies.

---

With the rapid growth of human population, food and nutrition shortage becomes a global challenge, and the utilization of unheeded or orphan crops potentially provides an alternative solution of this problem (Naluwairo 2011). *Lablab purpureus* (Dolichos bean) and *Vigna subterranea* (Bambara groundnut) are members of tribe Phaseoleae in Fabaceae, both of them are widely environmental adaptive and with the ability of nitrogen-fixing, regarded as important protein sources for African (Robotham and Chapman 2017; Gbaguidi et al. 2018). Here, we report the complete plastid genomes of the two neglected crops *L.purpureus* and *V. subterranea* on the basis of Illumina paired-end sequencing data.

In this study, fresh leaves were collected from plants that grow at the World AgroForestry Center (ICRAF) campus that are supported by the African Orphan Crops Consortium (AOCC). The total genomic DNA was extracted using a modified CTAB method (Sahu et al. 2012), and was used to construct the paired-end libraries with 250 bp insert size. Then the libraries were sequenced on a HiSeq 2000 platform (Illumina, San Diego, CA), and 47 and 36 Gb high quality reads were generated for *L. purpureus,* and *V. subterranean,* respectively. SOAPfilter v2.2 was used to trim the raw reads with the following criteria (1) reads with >10 percent base of N; (2) reads with >40 percent of low quality (quality value ≤10) sites; (3) reads contaminated by adaptor sequence or produced by PCR duplication. Finally, 40 Gb clean data of *L. purpureus* and 33 Gb clean data of *V. subterranea* were kept. Plastid genome was assembled based on NOVOPlasty v2.5.9 (Dierckxsens et al. 2016) using *Arabidopsis thaliana* plastid genome (NC_000932.1) as a seed. The plastome of *Phaseolus vulgaris* (DQ886273.1) and *Vigna radiata* (NC_013843.1) were used as references for the contig extending of *L. purpureus* and *V. subterranea* using the program MITO bim v1.8 (Hahn et al. 2013). The annotation of protein-coding genes were conducted with GeneWise v2.4.1 (Birney et al. 2000), and further verified on the web application GeSeq (Tillich et al. 2017). The annotation of transfer RNA (tRNA) and ribosomal RNA (rRNA) genes were performed using the program GeSeq (Tillich et al. 2017). The complete plastid genomes of *L. purpureus* and *V. subterranea* have been submitted to CNGB Nucleotide Sequence Archive (CNSA: https://db.cngb.org/cnsa; accession number: CNA0000811 and CNA0000812).

The complete plastid genome of *L. purpureus* and *V. subterranea* have a typical quadripartite structure, containing two inverted repeats (IRs), a large single-copy region (LSC), and a small single-copy region (SSC). *L. purpureus* has a total length of 151,753 bp, with two 26,054 bp IRs, a 81,727 bp LSC and a 17,918 bp SSC, while *V. subterranea* plasid genome is 152,015 bp long, with two 26,159 bp IRs, a 82,157 bp LSC and a 17,540 bp SSC. For the gene content, both plastomes encode 71 protein-coding genes and four rRNA genes, *L. purpureus* has 32 tRNA genes, while *V. subterranea* has 33 tRNA genes. In both plastomes, twelve genes (*rps19*, *rpl2*, *rpl23*, *ycf2*, *ndhB*, *rps7*, *rps12*, *trn*N-GUU, *trn*R-ACG, *trn*A-UGC, *trn*V-GAC, *trn*L-CAA and *trn*M-CAU) are duplicated in the IR regions.

To reconstruct the phylogenetic relationships of the two orphan crops and other Fabaceae species, 17 published plastomes from tribes of Phaseoleae and Millettieae of Fabaceae were retrieved from the NCBI database. Seventy-seven protein-coding genes were extracted from all samples, and used for gene alignment using the program MAFFT v7.017 (Katoh et al. 2013). The aligned genes were concatenated into a data matrix of 67,020 sites. The Maximum-Likelihood tree was constructed using this concatenated dataset under the GTRCAT model using RAxML v8.2.4 (Stamatakis 2014) with 100 bootstraps. The phylogenetic analysis showed that *L. purpureus* and *Phaseolus vulgaris* were closely related, and *V. subterranea* and other *Vigna* species are clustered in the same clade (Figure 1), which is congruent with former studies (Wang et al. 2017).

**Figure 1.**
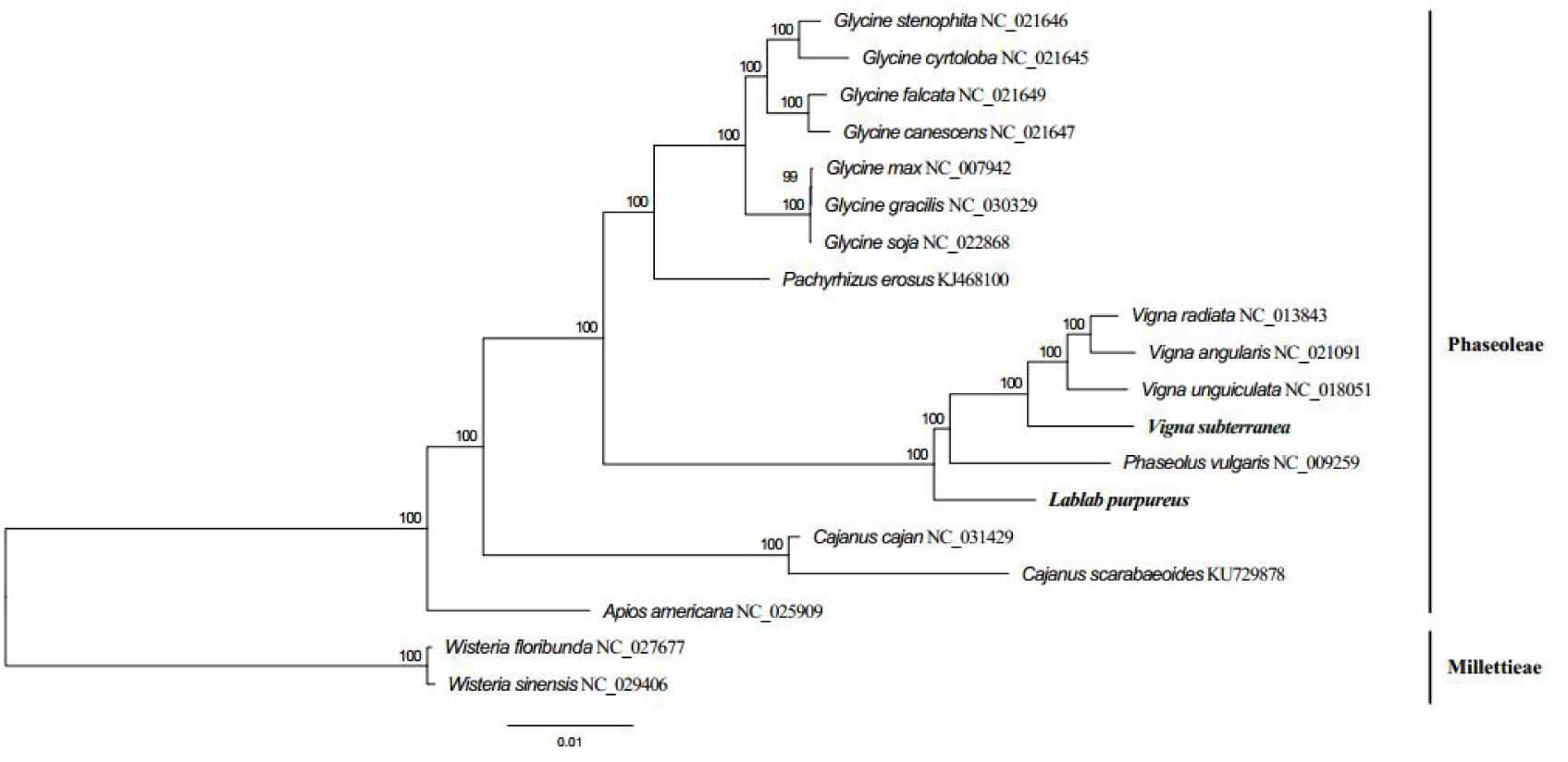
Maximum-likelihood tree based on the nucleotide sequences of 77 plastid protein-coding genes from 19 plastid genomes of Phaseoleae and Millettieae of Fabaceae.

## Disclosure statement

No potential conflict of interest was reported by the authors.

## Funding

This work was supported by the grants of Basic Research Program, the Shenzhen Municipal Government, China (No.JCYJ20150529150409546) and (No.JCYJ20150831201643396), as well as the funding to State Key Laboratory of Agricultural Genomics (No.2011DQ782025), and Guangdong Provincial Key Laboratory of Genome Read and Write (No.2017B030301011).

